# Three-dimensional total-internal reflection fluorescence nanoscopy with nanometric axial resolution by photometric localization of single molecules

**DOI:** 10.1101/693994

**Authors:** Alan M. Szalai, Bruno Siarry, Jerónimo Lukin, David J. Williamson, Nicolás Unsain, Damián Refojo, Alfredo Cáceres, Mauricio Pilo-Pais, Guillermo Acuna, Dylan M. Owen, Sabrina Simoncelli, Fernando D. Stefani

## Abstract

Single-molecule localization microscopy (SMLM) enables far-field imaging with lateral resolution in the range of 10 to 20 nanometres, exploiting the fact that the centre position of a single molecule’s image can be determined with much higher accuracy than the size of that image itself. However, attaining the same level of resolution in the axial (third) dimension remains challenging. Here, we present SIMPLER, a photometric method to decode the axial position of single molecules in a total internal reflection fluorescence (TIRF) microscope. SIMPLER requires no hardware modification whatsoever to a conventional TIRF microscope, and complements any 2D SMLM method to deliver 3D images with nearly isotropic nanometric resolution. Examples of the performance of SIMPLER include the visualization of the nuclear pore complex through dSTORM with sub-20 nm resolution and of microtubule cross-sections resolved with sub-10 nm through DNA-PAINT.

## Introduction

Imaging the three-dimensional organization of biological structures down to the size of their structural proteins, ~ 4 to 10 nm, can open up exciting opportunities in the life sciences. Super-resolution microscopy, also known as far-field fluorescence nanoscopy, has set the conceptual pathway to achieve this goal^1–6^. Whereas in theory all super-resolution methods are able to reach nanometric resolution given a sufficiently high fluorescence photons flux, in practice most methods reach a lateral resolution limit of 10 to 20 nm. Axial resolution of methods using a single objective lens is typically two to five fold worse^7,8^, including recent advances considering the experimentally determined microscope point-spread-functions^9^, intensity-based approaches that rely on supercritical angle fluorescence^10^ or photometric analysis of the defocused images of single molecules^11^. By exploiting the 4Pi configuration^12^ it is possible to reach an axial resolution below 35 nm, but at the cost of increased technical complexity. Isotropic STED (isoSTED) has been shown to deliver nearly isotropic resolution in the range of 30 to 40 nm^13,14^, whereas 4-Pi PALM/STORM has reached 10 to 20 nm resolution in 3D^15–17^. To date, sub-10 nm axial localization of single molecules was only achieved by two methods, MIET and MINFLUX. Metal induced energy transfer (MIET) decodes the *z*-position of fluorophores through lifetime imaging making use of the distance-dependent energy transfer from excited fluorophores to a metal film^18^ or a graphene sheet^19,20^. However, combining this nanosecond time-resolved method with other nanoscopy methods in order to obtain 3D imaging with sub-10 nm resolution is not straightforward^21^. More recently, MINFLUX^24^ was demonstrated to deliver sub-10 nm resolution in three dimensions^23^, but this is at the cost of elevated technical complexity.

The use of the evanescent illumination field of total internal reflection to obtain sub-diffraction axial information in optical microscopy goes back to the 1950s and 1960s^24,25^. TIRF microscopy was pioneered by Axelrod in the early 1980s, demonstrating various applications including a scheme to obtain sub-diffraction axial resolution analysing the TIRF intensity as a function of the incidence angle^26^. These initial approaches were based exclusively on the axial dependency of the excitation intensity. Lanni et al. were the first ones to obtain axial positions from photometric readings of TIRF-illuminated 3T3 fibroblast cells excited at two different angles of incidence, and a theoretical calibration based on the model of Lukosz^27–29^. They could estimate average cell-substrate separation distances of 49 nm for focal contacts, and of 69 nm for close contacts^30^. In 1987 Axelrod revisited the work of Lukosz in the context of TIRF microscopy^31^, and from then on, the knowledge to obtain axial positions from a quantitative use of TIRF microscopy was fully available.

Here, we introduce an easy-to-implement photometric method named SIMPLER (Supercritical Illumination Microscopy Photometric *z*-Localization with Enhanced Resolution) to determine the axial position of single molecules in a total internal reflection fluorescence (TIRF) microscope. SIMPLER is able to deliver an axial localization accuracy comparable to MIET or MINFLUX, and at the same time requires no hardware modifications whatsoever to a conventional TIRF microscope and is fully compatible with all SMLM methods. We demonstrate the performance of SIMPLER in combination with DNA-PAINT and dSTORM, achieving nearly isotropic resolutions of 8 and 20 nm, respectively, throughout an axial range of 250 nm.

## Results

### Principle of SIMPLER and theoretical axial localization precision

Basically, all methods to obtain axial positions of molecules from TIRF measurements involve two parts, a calibration of the TIRF signal and a method to estimate the axial position. The calibration of the TIRF signal can be obtained theoretically or experimentally. Theoretical calibration of the TIRF signal implies knowing of the excitation evanescent field and the angular emission pattern of molecules at different *z*-positions. Calculating the evanescent field is a relatively straightforward task. By contrast, calculations of the angular emission pattern of molecules as a function of the distance to the interface are usually only at hand for someone with expertise in optics. Presumably for this reason, most works attempted to obtain calibrations using different experimental approaches. Examples include scanning axially quantum dots or fluorescent beads with a piezo actuator^32–34^, or analysis of the emission intensity of fluorescent species fixed at different separations from the substrate. For the latter strategy, fluorophores were attached to a convex lens^35,36^, large spherical beads^37^, tilted microtubules^38,39^, a tilted glass coverslip^40,41^, or a nanometric staircase structure^42^. Remarkably, in most of these works, the effect of the substrate-sample interface on the emission power and angular emission was not present in the conceptual discussion. We note that, although an experimental calibration of the TIRF signal could include this effect, it is necessary considering the effect of the extra interface used to hold the calibration fluorescent probes in place, which will influence the detected signal too. Only in exceptional cases this was taken into account by matching the refractive indices of the liquid and the holding material^37,42^, but this approach has a shortcoming too because the liquid used for the calibration usually has a refractive index different from the real samples.

With the calibration data at hand, previous methods have obtained axial positions or relative distances between fluorophores in two different ways. The one method estimates relative positions of particles within the evanescent field from the ratio of its fluorescence intensity under TIR and wide-field illumination^33,43,44^. The other method, usually called variable-angle TIRF, obtains absolute *z*-positions from TIRF measurements of the same sub-diffraction object at two^30^ or more incidence angles^39,45–50^, and fitting the intensity vs. incidence angle according to the calibration model. In order to determine the axial position of an object, both types of methods require sequential measurements of its emission under different illumination conditions. This is hardly compatible with the fast single molecule blinking required for SMLM. Two recent works clearly demonstrate this limitation. On the one hand, Jung et al. made correlative measurements of cell membrane topography using variable-angle TIRF microscopy and SMLM. They could determine the membrane topography with 10-20 nm axial resolution but with diffraction-limited lateral resolution, and vice-versa for T-cell receptors^51^. On the other, Fu et al.^52^ applied variable-angle TIRF to perform an effective optical sectioning with a axial resolution of 20 nm; each section was defined as the difference in the imaging depths obtained at the two adjacent angles of incidence. In each section, they performed SMLM and obtained a lateral resolution of about 100 nm.

By contrast, SIMPLER decodes *z* information directly from 2D SMLM data. Through a full theoretical modelling, including the evanescent illumination, the modulation of the angular emission and the shape of the single molecule signals in the image plane, we demonstrate that the TIRF intensity signal can be effectively represented by just three parameters, which are easily accessible. This parametrization allows the determination of the axial position of individual molecules from a single measurement of their emission intensity, which in turn enables the direct combination of SIMPLER with any SMLM method to obtain 3D super-resolved images.

Figure 1 illustrates the concept of SIMPLER. Total internal reflection occurs when light incides from a medium with refractive index *n*_*i*_ on an interface with another medium of smaller refractive index *n*_*s*_ < *n*_*i*_. If the angle of incidence *θ*_*i*_ is larger than the critical angle *θ*_*C*_ = arcsin(*n*_*s*_/*n*_*i*_), light is fully reflected at the interface and an evanescent field appears, penetrating the medium of low refractive index with an intensity that decays exponentially. In a fluorescence microscope, TIR illumination can be generated by controlling the angle of incidence of the excitation light using an immersion objective lens as schematically shown in Figure 1a. In practice, the excitation field contains also a non-evanescent component due to scattering in components of the optical system, that decays on a much longer scale^37^. Near the interface, the non-evanescent component can be considered constant and the overall illumination field is represented by a linear superposition of both contributions, *I*(*z*) = *αI*_0_*e*^−*z*/*d*^ + (1 − *α*)*I*_0_ with *I*_0_ the intensity at the interface, 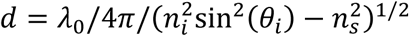 the penetration depth, *λ*_0_ the vacuum wavelength, and 1 − *α* the scattering contribution fraction. The excitation rate of a freely rotating fluorophore (under linear excitation) will depend on its axial position according to *I*(*z*). Figure 1b shows *I*(*z*) for one of our experimental configurations (*λ*_0_ = 642 nm, *n*_*i*_ = 1.517, *n*_*s*_ = 1.33 water, *θ*_*i*_ = 69.5°, *α* = 0.9), which decays with *d* = 102 nm.

**Figure 1.**
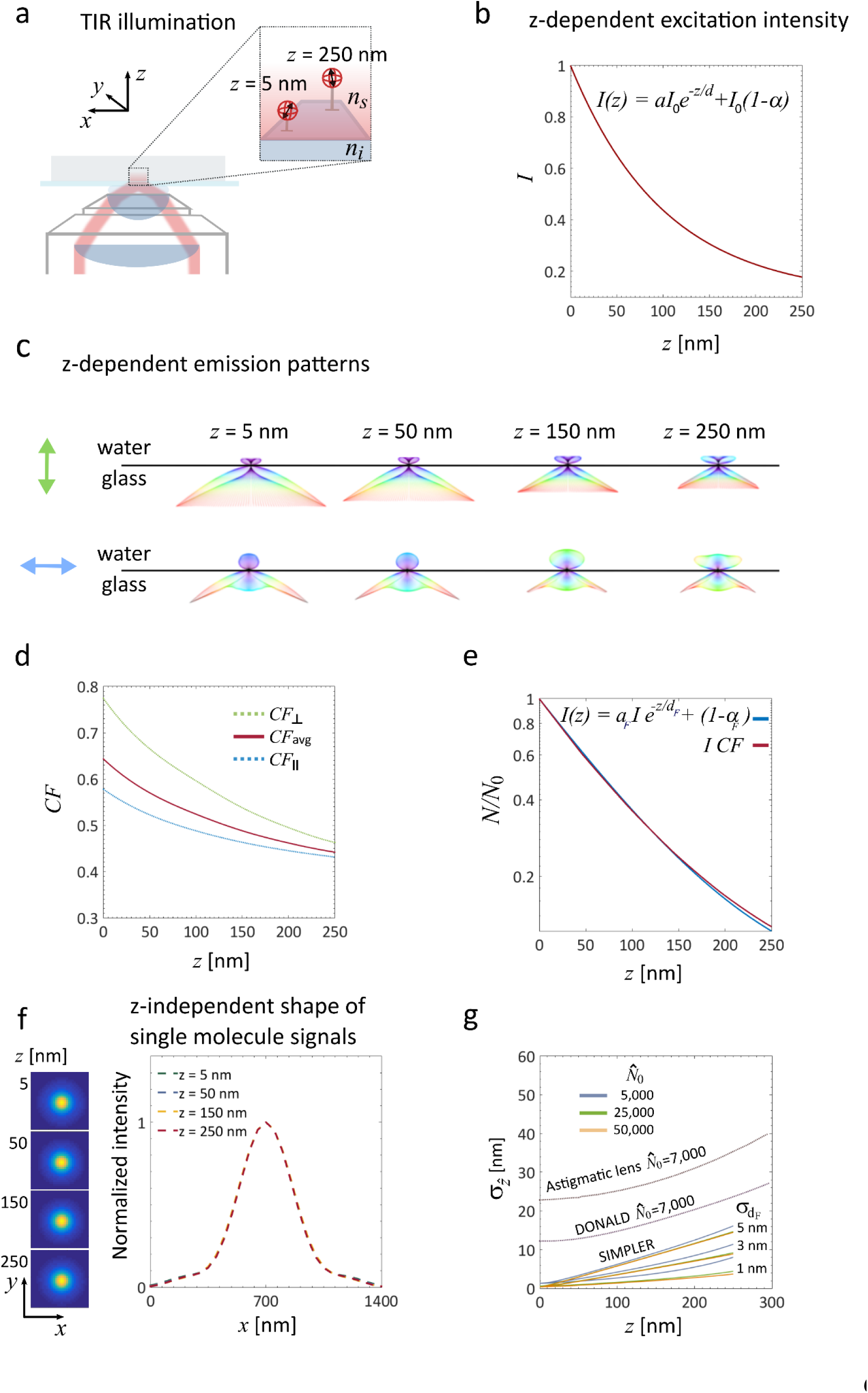
Supercritical Illumination Microscopy enables Photometric Localization Enhanced z-Resolution (SIMPLER). **(a)** Simplified optical layout of the excitation path of a TIRF microscope exemplifying isotropic emitters, which corresponds to the case of the dipolar emitter rotating faster than the measurement time. (**b**) Intensity of the excitation field under TIR illumination for our experimental configuration: *λ*_0_ = 642 nm, *n*_*i*_ = 1.517, *n*_*s*_ = 1.33 water, *θ*_*i*_ =69.5°, *α* = 0.9. (**c**) Simulated angular emission patterns of a dipolar emitter oriented either perpendicular (up) or parallel (bottom) to the water-glass interface and located at 5, 50, 150 and 250 nm above it. All calculations are made for *λ*_0_ = 670, the maximum emission wavelength of the fluorophores. (**d**) Fraction of fluorescence signal collected (collected fluorescence - CF) with the microscope objective (NA = 1.42) for a fluorophore emitting at 670 nm and oriented parallel (blue dotted line) or perpendicular (green dotted line) to the glass/water interface, normalized to the case of a molecule far from the interface. The solid red line represents the isotropic average corresponding to the case of a rotating fluorophore. (**e**) Calculated z-dependent fluorescence signal (represented in logarithmic scale) of single molecules expressed as the ratio of number of photons detected at a given *z* position (*N*) and the number of photons of an identical emitter placed at *z* = 0 (*N*_0_). Principle of SIMPLER: the axial position of single molecules is retrieved from *N*/*N*_0_ either through the exact solution (solid red line) or through the exponential approximation (solid blue line – equation 2). (**f**) Calculated images of single molecules at *z* = 5, 50, 150 and 250 nm with normalized intensity; *xy* images at the focal plane (right) and profiles along *x* (left). (**g**) Theoretical lower bound for the axial localization precision of SIMPLER for different sets of *N*_0_ and 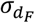. Comparison of the theoretical localization precision of SIMPLER with respect to the reported precision of two other well-stablished *z*-localization techniques: single lens astigmatism and DONALD, for 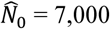 photons (data taken from^10^).

Within the range of TIRF, the process of fluorescence emission is also influenced by the dielectric interface^53^. Figure 1c shows the calculated angular emission patterns of fluorophores oriented parallel and perpendicular to the glass-water interface, for four different axial positions within the penetration depth of the evanescent field (*z* = 5, 50, 150 and 250 nm). Clearly, for both emitter orientations, fluorophores emit more fluorescence into the glass semi-space as they get closer to the interface. The dotted curves in Figure 1d are the integrals of the angular emission pattern over the collection solid angle of a microscope objective with *NA* = 1.42. These curves represent the collected fluorescence (*CF*) from single molecules oriented parallel and perpendicular to the interface, as a function of the axial position, and normalized to the case of a fluorophore far from the interface. In addition, the isotropic average (*CF*_*avg*_) is also shown, which corresponds to the usual experimental situation of rotating fluorophores.

Then, for a single molecule located in the evanescent field, the detected fluorescence signal will be proportional to the product of the excitation field and the collected fluorescence: *F*(*z*) = *I*(*z*) × *CF*_*avg*_(*z*) (hereafter referred to as the exact solution). As shown in Figure 1e, it turns out that *F*(*z*) is well represented by an exponential function analogous to *I*(*z*) but with a steeper decay (*d*_*F*_ = 87.5 nm) and smaller background constant (*α*_*F*_ = 0.93). The difference between the exact solution and the exponential approximation is negligible (< 1 nm for *z* < 150 nm, and < 8 nm for *z* < 250 nm) (Supplementary Figs. 1a and 2). This dependency of the fluorescence signal with the axial position is the core of SIMPLER axial localization.

In the context of SMLM, it is convenient to express the fluorescence signal in terms of the number of photons, *N*, detected in a given unit of time (typically the acquisition time of a camera frame):

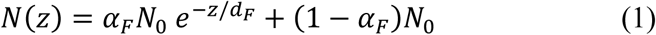

where *N*_0_ is the number of photons emitted by a fluorophore at *z* = 0.

Also relevant for SMLM is the fact that the variations of the angular emission emission pattern do not produce any significant modification of the shape of single molecule signals in the image plane. Figure 1f shows the calculated single molecule signals obtained by focusing the angular emission patterns of molecules at *z* = 5, 50, 150 and 250 nm, and normalized profiles. This means that *N*(*z*) can be estimated with a single procedure throughout the TIRF range.

Next, we analyse the theoretically maximum accuracy of SIMPLER for axial localization of single molecules using the exponential expression of *N*(*z*) as described in equation (1). Following equation 1, an experimental estimation of the axial position of a molecule 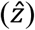 can be obtained from a measurement of the number photon counts detected in a camera frame time 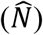, knowing the value of photon counts at 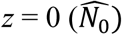 for an identical emitter:

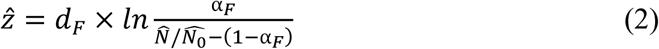

Then, standard error of 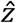, which ultimately determines the axial resolution in SMLM, can be estimated as:

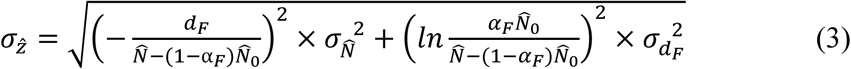

This expression is an approximation to illustrate the influence of the most important parameters and to obtain a theoretical lower bound for the axial localization error. It neglects the contributions of the uncertainties in 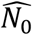 and *α*_*F*_. The reason is twofold. First, these two parameters are fixed for a given experiment (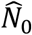 depends on the nature of the fluorophore and the experimental conditions, *α*_*F*_ is a fixed characteristic of the experimental set-up), and can, in principle, be determined with high accuracy in independent measurements. Second, the influence of their uncertainty is of minor importance, as it will be seen later. Under these conditions, for a given microscope/sample set-up (*i.e.* a given set of *d*_*F*_ and *α*_*F*_), 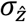 depends on the values of *N*_0_, 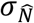 and 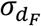. In order to compute a theoretical lower bound, we considered 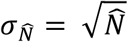 which arises from the fact that is Poisson distributed and that 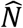 in SMLM the photon counts of each fluorophore are typically determined in one single measurement. We note, however, that in real life experiments, other factors may enlarge this value. For example, the variance introduced by EM amplification in EM-CCD cameras used in SMLM can lead to errors in photon counts that are a factor of 2 larger than Poisson statistics^54^. Figure 1g displays curves of 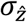 as a function of the axial position for experimentally accessible values of *N*_0_ and 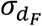, showing that SIMPLER is potentially able to deliver axial resolutions of a few nanometres under usual experimental conditions. Interestingly, a useful range with axial resolution below 10 nm can be obtained, depending on the uncertainty of *d*_*F*_. For 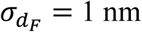, the sub-10 nm axial resolution range extends up to *z* = 250 nm, for 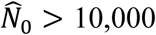. If 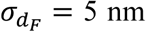 or larger, the resolution becomes fairly independent of the photon count for 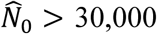, but the range of sub-10 nm resolution is limited to *z* < 170 nm. For comparison, Figure 1f also shows the performance of two other methods for axial localization^10^. Within the TIR penetration depth, SIMPLER is in principle able to achieve superior performance than the commonly used single cylindrical lens configuration^7^ and more recent approaches that decode axial position from fluorescence emission at supercritical angles (*i.e.* DONALD^10^ and SALM^55^). At the same time, SIMPLER holds the additional advantage that it does not require any hardware modification whatsoever to a conventional SMLM TIRF microscope, and that data is acquired and analysed in essentially the same way.

### Experimental implementation of SIMPLER

We performed experiments to characterize SIMPLER in two different SMLM TIRF microscopes. A custom-built microscope and a commercial microscope with flat-illumination optics and an internal calibration of the angle of incidence (Nikon N-STORM 5.0). Further details of each set-up are provided in the Methods section.

For SIMPLER, data is acquired and analyzed as in any other 2D SMLM method. The only difference is that for each single molecule emission event, in addition to the *xy* location, the number of photons per frame 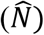 must be determined in order to obtain the molecular *z*-coordinate through equation (2). We note that most of the available software for SMLM image reconstructions already count with routines to output 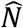.

The acquisition conditions should be adapted to record the average single molecule emission event during more than three camera frames. This enables an additional filtering step during analysis to exclude the first and last frames of each single molecule emission event for the determination of 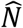 (Figure 2a), as it is uncertain whether the molecule was emitting or not during the complete integration time of those frames. This post-processing step in the reconstruction of 3D images is necessary to rule out low-intensity events that would bias axial localizations to artificially higher *z*-values.

**Figure 2.**
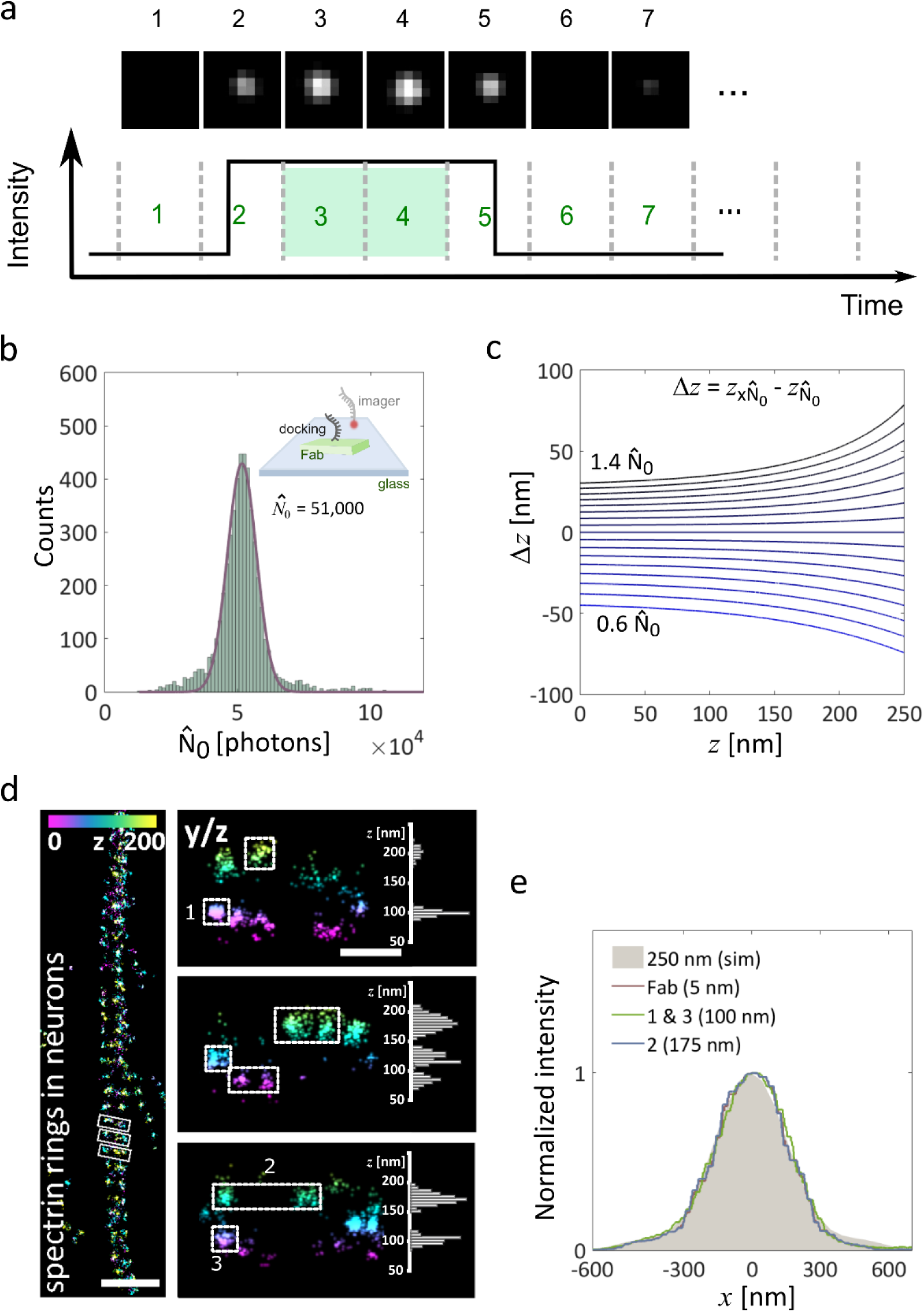
Data analysis and *z* assessment via SIMPLER. **(a)** Single molecules are detected and localized using any single molecule localization microscopy software. In the case of non-uniform illumination experiments (*i.e.* due to Gaussian shape of the excitation beam), the number of photons of each localization is corrected to the local excitation intensity (see Methods). Next, a frame filtering step is performed to only use localizations that lasted at least three frames and compute their photon count per frame excluding potentially misleading first and last frames. (**b**) Histograms of photons-counts for ATTO655, imaged using DNA-PAINT, with sample of DNA-labelled (docking strand) Fab fragments deposited over a coverslip. Imaging area was 16 x 16 µm^2^ (see Methods). Inset: Schematic representation of the experimental conditions. (**c**) Systematic error in *z*-localization (Δ*z*) as a function of *z* due to wrong values of *N*_0_ ranging from 0.6*N*_0_ to 1.4*N*_0_ in 0.05*N*_0_ steps. (**d**) β2-spectrin rings in hippocampal neurons. Left: top view (*xy*). Right: magnified side-views (*yz*) of the boxed regions in the top view, together with axial profiles of the boxed areas. Localizations are color-coded according to their *z*-position. (**e**) Normalized image profiles of single emitters located at different axial positions (*z* = 5, 100 and 175 nm) within the TIRF region along with the simulated profile for a rotating dipole located at *z* = 250 nm. Experimental profiles were obtained from samples of Fab fragments adsorbed to the coverslip (*z* = 5 nm) and β2-spectrin rings in hippocampal neurons (*z* =100 and 175 nm, numbered regions in **d**).

Determination of *z*-coordinates through equation (2) requires previous knowledge of *N*_0_, *d*_*F*_ and *α*_*F*_. Next we explain how these parameters are obtained.

*N*_0_ is the number of detected fluorescence photons for fluorophores located at *z* = 0. Thus, it can be determined by computing the average 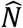 for fluorophores bound to structures whose distance to the coverslip is negligible. Such structures could be present in the same biological sample (*e.g.* if a known cellular component is known to be attached or very close to the substrate), or in another sample made specially to obtain *N*_0_. In the latter case, we found it practical to deposit directly on a coverslip the same fluorescent labels used for biological imaging and determine their average 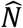 under identical experimental conditions as in the biological imaging experiments. For example, for DNA-PAINT experiments we simply deposited on the coverslip the same DNA-coupled secondary antibodies fragments (Fab) used for labelling the biological samples, and imaged them under identical conditions with the complementary fluorescently labelled DNA imager strand (Figure 2b, inset).

Far from saturation, *N*_0_ depends proportionally on the excitation light intensity, which in general is not uniform throughout the complete field of view of a wide-field microscope. In most cases, wide-field illumination is achieved using the central part of an expanded (nearly Gaussian) beam. Alternatively, some microscopes include special optics, such as apodizing neutral filters, to attain a flat(ter) illumination profile. We have tested SIMPLER in both kinds of set-up. In both cases, local variations of excitation intensity maybe also be present due to multiple, accumulative effects through the optical system. Thus, the photon count of each molecule used to obtain *N*_0_ was corrected using the local background level as a measure proportional to the local excitation intensity (see Methods for details about the correction procedure). Figure 2b shows a histogram of corrected photon-counts from fluorophores distributed over the field of view of our custom microscope (16 x 16 μm^2^), illuminated with the central part of an expanded Gaussian beam. In this case, the excitation intensity at the periphery of the field of view was 25% lower than in the centre. The corrected histogram of 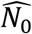 is well described by a normal distribution with an average value of 51,000 photons and a standard deviation of 10%. It turns out that a variability in 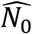 of ± 10%, mainly produces an off-set in the determined z-coordinates that ranges from ± 8 nm (for *z* = 0) to ± 18 nm (for *z* = 250 nm) (Figure 2c). Hence, imaging with this level of uncertainty in 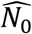 leads to average accumulated distortions smaller than 10 nm within the 0 - 250 nm axial range. It should be noted that this small variable error in *z*-localization is distributed throughout the field of view of 16 x 16 μm^2^. Considerably narrower distributions of 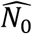 and smaller variations in *z*-localization are observed in smaller fields of view (see Supplementary Figure 3). This effect is even smaller in microscopes with flat illumination optics, and it becomes negligible 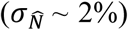 when analysing nanostructures located at almost fixed *xy* position.

**Figure 3.**
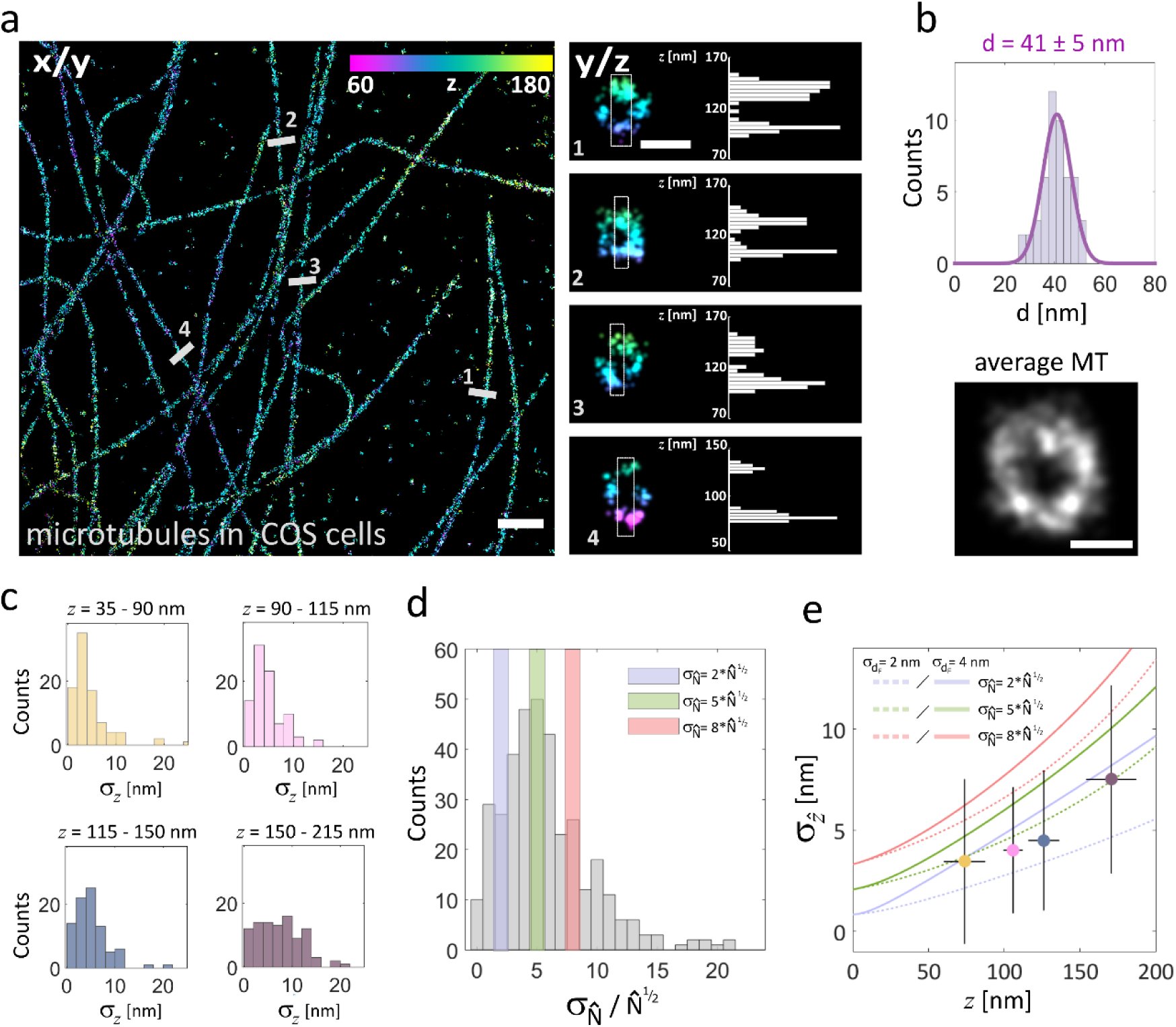
Microtubules immunolabeled for DNA-PAINT super-resolved in 3D using SIMPLER. (**a**) Microtubules in COS-7 cells. Left: top view. Right: magnified side-views along the numbered lines in the top view, together with the axial profile of the boxed areas. (**b**) Distribution of microtubule diameters (*n* = 50), with an average of 41 nm and a standard deviation of 6 nm. An average (*n* = 8) microtubule profile is also shown. (**c**) Histograms of 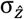 at different *z* positions, obtained experimentally from 381 DNA-PAINT single molecule traces. Data includes traces from microtubules in COS-7 cells, as well as from spectrin in hippocampal neurons (Figure 2d). (**d**) Corresponding 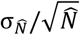 histogram of the same experimental data as in (**c**). (**e**) Median 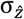 values located in the median *z*-position of each interval, overlapped with theoretical curves of 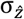 for 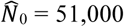 and 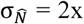, 5x or 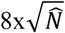 for σ_dF_ = 2 and 4 nm. Error bars represent the standard deviation of 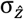 and *z*. Scale bars represent 1 µm (**a**, top view); 50 nm (**a**, side view) and 25 nm (**b**).

Next, we explain how to obtain *d*_*F*_ and *α*_*F*_ and how to optimize their values through the evaluation of 3D SIMPLER SMLM images. As explained in the previous section, *d*_*F*_ and *α*_*F*_ are obtained from a fit to the expected *z*-dependent single molecule fluorescence intensity *F*(*z*) = *I*(*z*) × *CF*_*avg*_(*z*) (Figure 1e).

The evanescent component of *I*(*z*) requires the calculation of the decay constant 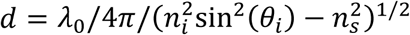, which depends on available experimental parameters. In the custom-built microscope, the angle of incidence was determined by analysing the lateral displacement of the focus as a function of axial displacement, as described in^42^ (Supplementary Figure 4). In the commercial microscope, we trusted the internal calibration of incidence angle.

**Figure 4.**
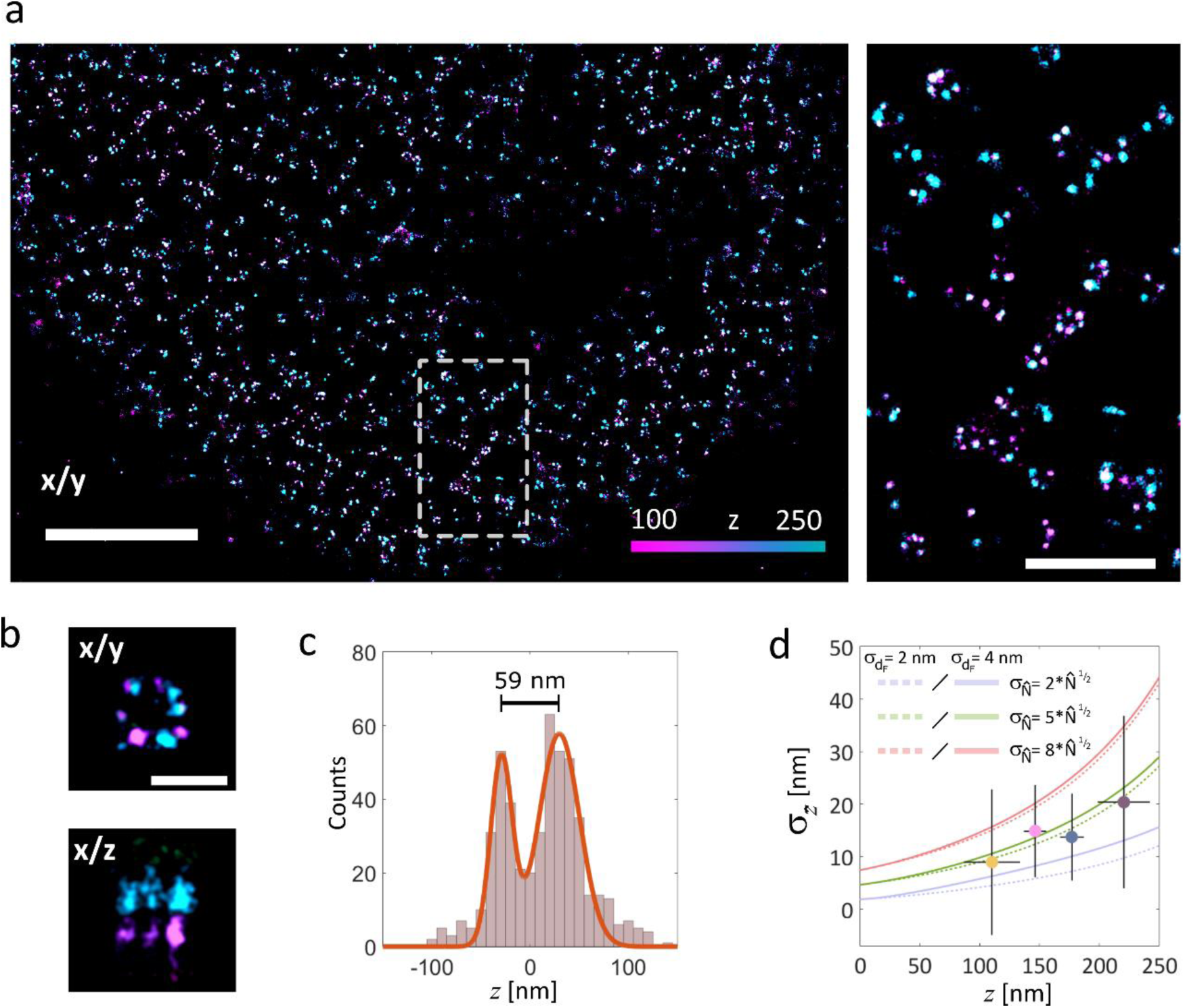
NUP107 immunolabeled for dSTORM and super-resolved in 3D using SIMPLER. (**a**) Left: top view of dSTORM image of NUP107-mEGFP in HeLa Kyoto cells, labelled with primary anti-GFP antibody conjugated to Alexa Fluor 647. Right: a magnified view of region of the region marked. (**b**) Average nuclear pore complex showing the archetypal eightfold symmetry (*xy* top-view, left) and the organization in nuclear and cytoplasmic rings (*xz* and *yz* side-views, center and right, respectively) reconstructed by SIMPLER (*n* = 4). (**c**) Histogram of *z* positions of the nuclear pore complex shown in (c) yields 59 nm separation between the nuclear and cytoplasmic rings. (**d**) Median 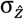 values of different *z* positions, obtained experimentally from 331 dSTORM single molecule traces overlapped with theoretical curves of σ_z_ for 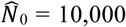 and 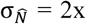, 5x or 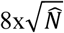 for σ_dF_ = 2 and 4 nm. Error bars represent the standard deviation of 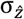 and *z*. Scale bars represent 2 µm (**a**, left); 500 nm (**a**, right) and 100 nm (**b**).

Determining accurately the relative contribution of the non-evanescent field (1 − *α*) can be challenging, as recently reviewed^56^. In objective-type TIRF microscopes (1 − *α*) is typically found to be between 10 to 15%^42,56^. Fortunately, SIMPLER is rather insensitive to variations *α* of in that range. For example, a misleading value of (1 − *α*) in the range of 8 to 12% introduces a maximum accumulated distortion smaller than 5 nm in the *z* range from 0 to 150 nm. This means that a structure occupying the axial range of 0-150 nm, would be imaged with an accumulated distortion smaller than 5 nm. For a larger structure, occupying the range from 0 to 250 nm, the total accumulated axial distortion would be of about 10 nm (Supplementary Figures 1b and 2). It should be noted that these are total accumulated distortions smaller than 4%, hardly noticeable and probably surpassed by other factors such as the size or the position of the labels.

Obtaining *CF*_*avg*_(*z*) requires the calculation of the angular emission pattern of single molecules and their integration over the numerical aperture of the objective. Details about these calculations are provided in the Methods section. In order to simplify this task for new users of SIMPLER, in the Supporting Information we provide tabulated values of *CF*_*avg*_(*z*) for the most usual TIRF microscopy configurations – *i.e. NA*’s ranging from 1.40 to 1.49 and fluorophores’ maximum emission wavelength (*λ*) between 500 nm to 720 nm (Supplementary Table 1). We also make available a Matlab routine that directly outputs *d*_*F*_ and *α*_*F*_ for each user input experimental conditions: *NA*; *λ*_0_; *λ*; *n*_*i*_; *n*_*s*_ and *α* (Supplementary Software).

The final component of SIMPLER is the invariability of the shape of the single molecule images. This was corroborated experimentally using 3D SIMPLER – DNA-PAINT images of the regular arrangement of β2-spectrin in the membrane-associated periodic skeleton (MPS) of neuronal axons. Figure 2d shows a top view of the MPS where its characteristic period of 190 nm is clearly visible. The 3D imaging using SIMPLER allows to resolve the sub-membrane organization of β2-spectrin across the axon and to identify single-molecule signals corresponding to spectrin molecules positioned at different heights. When normalized, all signals were indistinguishable, independently of their axial position. As an example, Figure 2d shows the normalized profiles of average signals obtained at 5 nm (Fabs deposited on the glass substrate), 100 or 175 nm (spectrin), and the calculated signal at 250 nm obtained from focusing the calculated emission pattern. Thus, a single algorithm can be used to obtain the photon counts of molecules positioned throughout the TIRF range. Additionally, we used this biological structure to compare 3D SIMPLER performance when the first and last frames of each single molecule trace are ruled out. In Supplementary Figure 5, it can be seen that different clusters are enlarged in the axial direction towards higher *z*-values when those frames are unfiltered, thus confirming the importance of performing this post-processing step.

### 3D SIMPLER SMLM of biological samples

Figures 3 and 4 illustrate the performance of SIMPLER SMLM to deliver super-resolved 3D images of biological structures. Figure 3 shows, as an example, a 3D image of the microtubule network of COS-7 cells imaged through DNA-PAINT. Figure 3a includes a top (*xy*) view of the microtubule network alongside with four cross-sectional views of individual microtubules. SIMPLER can fully resolve the hollow circular structure of immunolabeled microtubules, one of the smallest structural supramolecular protein structures in biological cells. Fitting a circle to 50 cross-sections of microtubules retrieves an average diameter of 41 nm with a standard deviation of 6 nm (Figure 3b), in good agreement with what is expected for an immunolabeled microtubule (primary antibodies and Fab fragments from secondary antibodies). In comparison to other methods that have achieved this level of axial resolution^9,17^, all of which involve high technical complexity, SIMPLER delivers equivalent or better resolution using the hardware of a conventional TIRF microscope with no modifications.

We use these images to determine experimentally the axial resolution 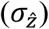 provided by SIMPLER – DNA-PAINT 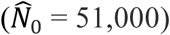 and compare it to the theoretical predictions. In order to obtain an experimental measure of 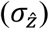, single molecule emission events longer than 5 camera frames were selected. In this way, after the first and last frame filter (Figure 2a), at least three independent measurements of 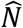, and their corresponding estimations of 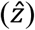, were available for each single molecule. Figure 3c shows the obtained distributions of experimental 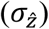 (366 single molecule traces) grouped for different ranges of *z*. The average resolution is well below 10 nm throughout the complete *z* range. Imaging with this level of resolution in 3D makes it possible to resolve bundles of microtubules that we found to be usual in hippocampal neurons, and that would otherwise be interpreted as single microtubules (Supplementary Figure 6). The hollow cross-sections of individual microtubules was also resolved through SIMPLER – DNA-PAINT images of microtubules in Human Fetal Foreskin Fibroblasts cells, acquired in a commercial TIRF microscope (Nikon N-STORM 5.0). In this case, we used the instrument internal calibration of the incidence angle, and applied SIMPLER directly, using the calculated parameters for that system and without even correcting for non-uniform illumination (the microscope was equipped with a gradient neutral-density filter to produce a near-flat-top intensity profile from the Gaussian-shaped beam input). These results (Supplementary Figure 7) demonstrate that SIMPLER is so robust that it can be applied to images acquired with commercial set-ups, making it available to any user who does not want to calibrate the incidence angles or correct for uneven illumination themselves.

The theoretical lower bounds of 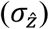 shown in Figure 1g were calculated using 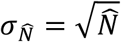, *i.e.* considering 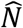 to be Poisson distributed. From the single molecule traces, we have determined the experimental variance of 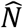 in single molecule emission traces and found larger variations. The distribution of experimental 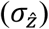 is shown in Figure 3d, which presents an average value of around 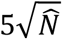. With this data at hand, we can make a direct comparison of the experimental performance of SIMPLER to the theoretical predictions. In Figure 3e, we plotted the median of the distributions of 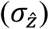 shown in Figure 3d against its *z* median. For comparison, we plotted the theoretical curves calculated from equation (3) for a set of values of 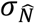 and 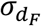. The experimental values of 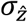 agree well with the theoretical prediction with values of 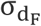 between 2-4 nm and 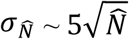

These results set boundaries to 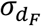 and 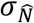. Therefore, the axial resolution achievable with SIMPLER in combination with any other SMLM method can be predicted using equation (3). As example, we provide 3D super-resolved images of the stereotypical arrangement of the nucleoporin Nup107 in the nuclear pore complex of HeLa Kyoto cells obtained via SIMPLER combined with dSTORM data, acquired in the Nikon N-STORM 5.0 microscope. Figure 4a shows the top-view images of a nucleus, where many nuclear pore complexes are visible. Even though the labelling efficiency was sub-optimal, the typical 8-fold symmetry of the complex is evident in many cases. More importantly, SIMPLER clearly resolves the axial separation of the cytoplasmic and nucleoplasmic rings. Figure 4b, shows lateral and axial cross-sections of an average nuclear pore complex. The axial separation distance between the cytoplasmic and nucleoplasmic rings is determined to be 59 nm, in excellent agreement with previous reports^57,58^.

The SIMPLER – dSTORM data was acquired with 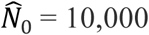 photons. A comparison of the experimental resolution (331 single molecule traces) to the theoretical prediction using 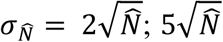 and 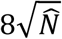 is shown in Figure 4d. As expected, the z-dependency of *σ*_*z*_ is in good agreement with the theoretical prediction with 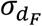 between 2 and 4 nm and 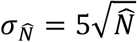. An axial resolution below 20 nm is achieved throughout the complete working range. With this level of axial resolution, the cross-sections of single microtubules and bundles (which are considerably smaller than the nuclear pores and thus far more challenging to visualize) are also visible. Supplementary Figure 8 shows example images of microtubule cross-sections obtained by 3D SIMPLER – dSTORM.

## Discussion

We have presented and characterized, theoretically and experimentally, a photometric method to localize single fluorescent molecules with nanometric precision in the axial direction of a total internal reflection fluorescence microscope. SIMPLER decodes the axial position (*z*) of single molecules based on three phenomena: the *z*-dependency of the excitation intensity, the *z*-dependency of the angular emission, and the *z*-independent shape of the single molecule signals in the image plane. A functional analysis of the *z*-dependent single molecule intensity enables its calibration based on just three parameters, that are easily accessible or provided in this work for most usual experimental configurations.

Because it delivers the axial position of molecules from a single intensity measurement, SIMPLER is fully compatible with any 2D SMLM. SIMPLER – dSTORM delivers 3D images with sub-20 nm axial resolution throughout the TIRF range of 250 nm, while for SIMPLER – DNA-PAINT the resolution is sub-10 nm. This level of axial resolution is only rivalled by methods of high technical complexity, such as 4-Pi nanoscopy, MINFLUX or MIET. By contrast, SIMPLER requires no hardware modification whatsoever to a wide-field single molecule fluorescence microscope and is highly robust; we validated its performance in custom-built microscopes and commercial instruments. Furthermore, unlike other 3D fluorescence nanoscopy methods, the level of resolution achieved by SIMPLER does not depend on nanometric axial drift corrections. This is because the measurement reference (the dielectric substrate-sample interface) is part of the sample. SIMPLER only requires axial stability provided by any standard focus-lock system (~ ± 20 nm over several hours).

In summary, making quantitative use of a TIRF microscope in combination with the concepts of super-resolution microscopy, it is possible to locate single molecules with nanometric accuracy simply from a measurement of its emission intensity. Due to its robustness and practicality, SIMPLER can be directly applied by any lab counting with a conventional TIRF SMLM microscope, making 3D fluorescence nanoscopy readily available to numerous users and enabling a new wave of discoveries about the structure and pathways of sub-cellular structures and protein-protein interactions.

## Methods

### Simulation of single molecule emission

The angular emission pattern of single molecules was calculated using a Finite Difference Time Domain solver (CST Microwave Studio). The molecules were considered as a small (1 nm) dipole oscillating at the frequency of emission. The fraction of detected fluorescence was obtained by integrating the emission pattern over the solid angle of interest. The single molecule images were obtained by focusing the fraction of the emission pattern collected by the objective. We provide sets of calculations for the most usual configurations in the Supporting Information, Supplementary Table 1.

### Super-resolution microscopy setup 1

The microscope used for TIR fluorescence SMLM of Figures 2d and 3a and Supplementary Figures 2, 3, 5, 6 and 8 was built around a commercial inverted microscope stand Olympus IX73 equipped with a high numerical aperture oil-immersion objective lens (Olympus PlanApo 60x / NA 1.42). Excitation was carried out with a circularly polarized 642 nm 1.5 W laser (MPB Communications 2RU-VFL-P-1500-642). TIR illumination was achieved with a linear translation stage (Thorlabs MT1-Z8) used to control the lateral position of the focused excitation beam on the back focal plane of the objective. The angle of incidence was set to 69.5° (Supplementary Method 1 and Supplementary Fig. 4). A dichroic mirror (Semrock Di03-R 405/488/532/635-t1) and a band-pass filter (Chroma ET700/75m) were used to separate the fluorescence emission of the sample from the laser excitation. The emission light was expanded with a 2x telescope so that the pixel size of the EMCCD camera (Andor iXon3 897 DU-897D-CS0-#BV) would match the optimal value for single-molecule localization (133 nm in the focal plane). The camera and laser were controlled with custom software developed in the laboratory and described in an earlier publication^59^. Typically, we acquired sequences of 50,000-100,000 frames at 4 Hz acquisition rate with a laser power density of ~2.5 kW/cm^2^ for DNA-PAINT, and 50 Hz, and ~ 3 kW/cm^2^ for dSTORM.

### Super-resolution microscopy setup 2

To demonstrate the ease of use of the technique SIMPLER was directly apply to 2D SMLM images acquired with a commercial Nikon N-STORM 5.0 system located in the Nikon Imaging Centre at King’s College London, UK and using the instrument internal calibration of the incidence angle (Figure 4a and Supplementary Fig. 7). The microscope is equipped with a 100× 1.49 numerical aperture oil immersion TIRF objective, a perfect focus system for stable axial drift-free imaging, a gradient neutral-density filter to produce a near-flat-top intensity profile from the Gaussian-shaped beam input and a CMOS camera (Orca Flash 4.0 V3, Hamamatsu). Samples were imaged under TIRF illumination with a 647 nm laser line that was coupled into the microscope objective using a quad band set for TIRF (Chroma 89902-ET-405/488/561/647 nm). The final pixel size of the image was 160 nm in the focal plane. We acquired sequences of 50,000-100,000 frames at 10 Hz acquisition rate with a laser power density of ~ 2.5 kW/cm^2^ for DNA-PAINT, and 50 Hz, and ~ 4 kW/cm^2^ for dSTORM using an angle of incidence of 69º.

### Primary neuron culture and cell lines

Mouse (CD1) hippocampal neurons were harvested from embryonic day 17 pups, following the general guidelines of the National Institute of Health (NIH, USA) and approval of the National Department of Animal Care and Health (SENASA, Argentina), and cultured in Neurobasal medium (Gibco) supplemented with 5 mM GlutaMAX-I (Gibco) and 2% B27 supplement (Gibco) at 37 °C and 5% CO_2_. Neurons were seeded at a density of 125 cells/mm^2^ on #1.5 thickness glass-bottomed chamber slides (Lab-Tek II, Thermo Fisher Scientific) and incubated for either 3 or 28 days, respectively. To increase cell attachment, glass slides were previously coated with 0,05 mg/mL poly-L-lysine (overnight at 37°C) (Sigma Aldrich) and 1 μg/μL Laminin (3 h at 37°C) (Sigma Aldrich).

Culture of COS-7, Human Fetal Foreskin Fibroblasts HFFF2 (ECACC 86031405) and HeLa Kyoto with endogenous Nup107 tagged with mEGFP (CLS Cell Lines Service GmbH) cell lines were grown in Dulbecco’s modified Eagle’s medium (DMEM) supplemented with 10% fetal bovine serum and 2 mM L-glutamine (Gibco) at 37 °C and 5% CO_2_.

### Sample fixation and permeabilization

Neurons, COS-7 and HFFF2 cells were fixed and permeabilized in PHEM buffer (60 mM PIPES, 25 mM HEPES, 5 mM EGTA, 1 mM MgCl2, pH=7.0), supplemented with 0.25% glutaraldehyde, 3.7% paraformaldehyde, 3.7% sucrose and 0.1% Triton X-100, for 20 min at room temperature. Auto-fluorescence was quenched by incubating the samples in 0.1 M glycine in PBS for 15 minutes followed by 3× washes with PBS. The fixed and quenched samples were blocked with 5% BSA in PBS containing 0.01% Triton X-100 for 1 h. HeLa Kyoto mEGFP-Nup107 cells were flash fixed and permeabilizing by sequentially using 2.4% paraformaldehyde in PBS for 30 sec and 0.1% Triton X-100 for 3 min. After 3× 5 min washes with PBS, cells were further fixed with 2.4% paraformaldehyde in PBS for 20 min and auto-fluorescence was quenched by incubating the samples in 0.1 M glycine in PBS for 5 minutes. followed by 3× washes with PBS.

### Immunostaining and imaging

Spectrin in neurons (Figure 2d and Supplementary Figures 2 and 5) was labelled with a mouse monoclonal primary antibody anti-β-Spectrin II (Clone 42/B-Spectrin II, BD Biosciences) for 1 h at room temperature using a 1:400 dilution in 5% BSA in PBS, followed by 3× washes with PBS. DNA-conjugated secondary antibody staining was performed by incubating the sample with a donkey anti-mouse secondary fragment antibody (Jackson ImmunoResearch, 715-007-003) at a 1:100 dilution in 5% BSA in PBS for 1 h at room temperature, followed by 3× washes with PBS. Microtubules in COS-7 cells (Figure 3a, Supplementary Fig. 2 and 8), HFFF2 cells (Supplementary Fig. 7) and in neurons (Supplementary Fig. 6 and 8) were treated with anti α-tubulin and anti β-tubulin primary antibodies for 1 h at room temperature using 1:400 dilutions in 5% BSA in PBS, followed by 3× washes with PBS (mouse monoclonal anti-α-Tubulin, clone TUB-A4A Sigma Aldrich; mouse monoclonal tyrosine anti-α-Tubulin, clone TUB-1A2 Sigma Aldrich; rabbit polyclonal anti-β-III-Tubulin, Abcam #ab 18207 for neurons; and rabbit polyclonal anti-β-II-Tubulin Abcam #ab 196 for COS-7 cells, kind gift of Dr. Jesus Avila, Centro de Biologia Molecular “Severo Ochoa” CBMSO, Consejo Superior de Investigaciones, Cientificas, Universidad Autonoma de MadridUAM, C/ Nicolas Cabrera, 1. Campus de Cantoblanco, 28049 Madrid, Spain60). In both cases secondary staining was done by 1 h treatment at room temperature with a mix of donkey anti-mouse DNA-conjugated secondary fragment antibody (Jackson ImmunoResearch, 715-007-003) and donkey anti-rabbit DNA-conjugated secondary fragment antibody (Jackson ImmunoResearch, 711-007-003) at a 1:100 dilution in 5% BSA in PBS for DNA-PAINT imaging or AlexaFluor647 conjugated goat anti-rabbit (Invitrogen, #A-21245) and AlexaFluor647 conjugated goat anti-mouse (Invitrogen, #A-21235) secondary antibodies at a 1:300 dilution in 5% BSA in PBS for dSTORM imaging, followed by 3x washes with PBS. Nup107 in HeLa Kyoto mEGFP-Nup107 cells were labelled with an Alexa Fluor 647 anti-GFP rat monoclonal primary antibody (Clone FM264G, BioLegend) for dSTORM imaging (Figure 4a) by incubating overnight at 4°C using a 1:200 dilution in 2% BSA in PBS, followed by 3× washes with PBS.

100 nm gold nanoparticles (BBI solutions) were added as fiducial markers for drift correction by incubating the sample for 5 min in a 1:2 solution of nanoparticles in PBS. After 3× washes with PBS, either PAINT buffer (Buffer B+: 5 mM Tris-HCl, 10 mM MgCl 2, 1 mM EDTA and 0.05 % Tween 20 at pH 8.0) containing fluorescently labeled DNA imager strands (Img1: ATTO655-5’-AGTTACATAC-3’ and Img2: ATTO655-5’-AGAAGTAATG-3’, biomers.net GmbH, for imaging anti-mouse and anti-rabbit DNA-conjugated secondary antibodies respectively) or STORM imaging buffer (IB) containing 50 mM TRIS pH 8, 10 mM NaCl, 10% w/v D-glucose, 10 mM mercaptoethylamine, 1 mg/mL glucose oxidase, and 40 μg/mL catalase were added to the immunolabeled samples.

Antibody conjugation to DNA-PAINT docking sites (5’-TATGTAACTTT-3’-Thiol and 5’-ATTACTTCTTT-3’-Thiol, biomers.net GmbH, for the donkey anti-mouse and the donkey anti-rabbit conjugates respectively) was performed using maleimidePEG2-succinimidyl ester coupling reaction according to a published protocol^61^ as described in Supplementary Method 2. Imager strands concentrations were 20 nM Img1 for spectrin imaging (Fig. 2d, Supplementary Fig. 2 and 5), and 80 pM Img1 + 80 pM Img2 for α-tubulin + β-tubulin imaging (Figure 3a, Supplementary Figures 2, 6 and 7). Samples were then used immediately for DNA-PAINT imaging.

### Data acquisition, analysis and 3D image rendering

Lateral (*x*,*y*) molecular coordinates and photon counts 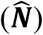 were obtained using the Localize module of Picasso software^61^ and enabling the symmetric PSF fitting method. Drift correction was carried out with a combination of redundant cross-correlation and fiducial markers approach using the Render module of Picasso. Photon-counts were corrected either by using the corresponding background parameter obtained from the MLE analysis or the illumination profile of the beam (measured by imaging a 1 μM Alexa Fluor 647 solution with the same incident angle as in the biological experiments).

Next, localizations were filtered to discard the frames corresponding to the switching (ON or OFF) of the fluorophores during the frame acquisition, whose photon count would be lower and lead to falsely high *z* coordinates. To ensure the molecule was emitting during the whole exposure time, localizations were kept as valid only in the case that other localizations, reasonably attributed to the same fluorophore (within a 3***σ***_***x,y***_ distance), were detected in the previous and subsequent frames (Figure 2a). Molecules detected for less than three frames were thus ignored.

For each image, a photon count was assigned to *z* = 0 (***N***_**0**_). This value was obtained directly by measuring samples of Fab fragments or antibodies adsorbed to the coverslip. For each localization, *z*-localization precision (σ_z_) was calculated as described in the main text (Figures 3e and 4e). Finally, z-color-coded image rendering was done using the ImageJ plug-in ThunderStorm^62^, importing the list of (*x, y, z*). A Gaussian filter with σ = 2 nm was used for all three dimensions. A lenient density filter was applied to *xy* images, to discard localizations with less than 100 neighbours in a 67nm radius, to enhance contrast by suppressing some of the non-specific localizations of the background.

### Data availability

The data sets generated and analyzed in this study are available from the corresponding author upon reasonable request.

## Supporting information

Supplementary Software

## Acknowledgements

This work has been supported by: CONICET, ANCYPT projects PICT2013-0792 and PICT-2014-0739, the Royal Society project IEC\R2\181018, the BBSRC grant BB/R007365/1, FOCEM (Fondo para la Convergencia Estructural del Mercosur) grant COF 03/11, and Swiss National Science Foundation through the National Center of Competence in Research Bio-Inspired Materials. S.S. acknowledges financial support from the Human Frontier Science Program Organization and the Royal Society through a HFSP and Dorothy Hodgkin fellowship, respectively. F.D.S thanks the support of the Max-Planck-Society and the Alexander von Humboldt Foundation. D.R. acknowledges the support of the Max Planck Society and the Volkswagen Stiftung.

## Author contributions

S.S. and F.D.S. conceived the approach and wrote the manuscript with input from all authors. A.M.S., B.S., S.S. and F.D.S. developed the analysis method. A.M.S., B.S. and S.S. acquired and analyzed the data. D.J.W. helped with the data analysis. A.M.S. and S.S. produced the custom-made DNA-PAINT antibodies. J.L, N.U., D.R. and A.O.C. contributed the primary neuron cultures and COS-7 cell lines, and carried out the spectrin and microtubules immunostaining. F.D.S., M.P-P and G.A. simulated the emission pattern of single molecules. D.M.O. discussed results and commented on the manuscript.

