## Supplementary Information for "Three-dimensional total-internal reflection fluorescence nanoscopy with nanometric axial resolution by photometric localization of single molecules"

| | |
|---|---|
| **Supplementary Figure 1** | Quantification of the differences in *z* values obtained using the exact solution of the fluorescence signal or the exponential approximation and the effect of varying $\alpha$. |
| **Supplementary Figure 2** | Comparison of side-view reconstructions by SIMPLER using different computation methods and varying $\alpha$ and $N_0$. |
| **Supplementary Figure 3** | Characterization of the uncertainty in z due to the correction of N by the excitation profile. |
| **Supplementary Figure 4** | Calibration of the TIRF excitation angle. |
| **Supplementary Figure 5** | Influence of the first and last frame filtering step on image quality. |
| **Supplementary Figure 6** | Microtubules from hippocampal neurons immunolabeled for DNA-PAINT super-resolved in 3D using SIMPLER in a custom-made setup |
| **Supplementary Figure 7** | Microtubules from Human Fetal Foreskin Fibroblasts cells, immunolabeled for DNA-PAINT super-resolved in 3D using SIMPLER using a commercial setup (Nikon STORM 5.0). |
| **Supplementary Figure 8** | Microtubules immunolabeled for dSTORM super-resolved in 3D using SIMPLER |
| **Supplementary Table 1** | Axial dependence of the collected fluorescence signal. |
| **Supplementary Method 1** | TIRF angle calibration |
| **Supplementary Method 2** | DNA-antibody coupling reaction |

### Supplementary Figures

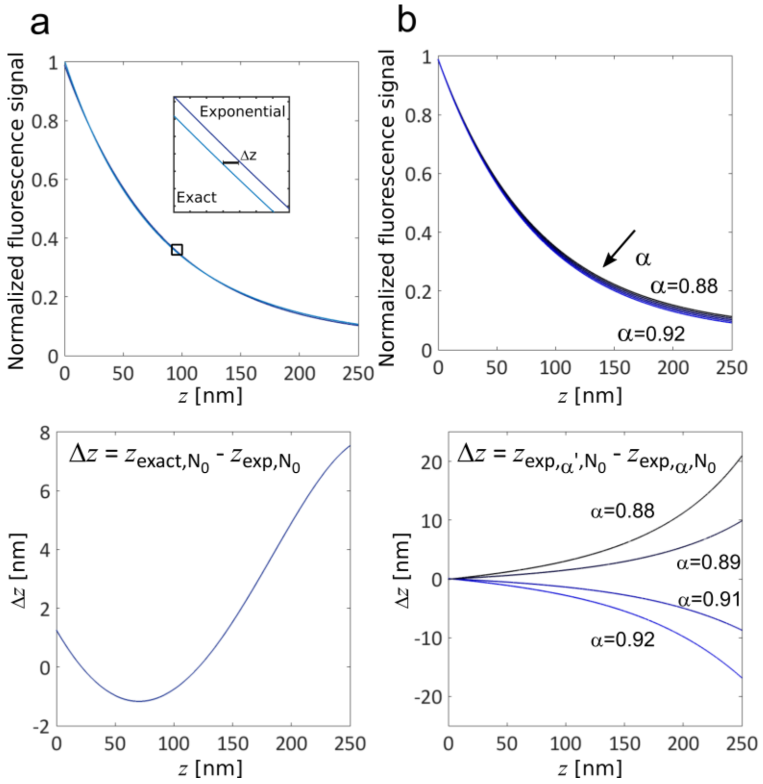

**Supplementary Figure 1. Quantification of the differences in *z* values obtained using the exact solution of the fluorescence signal or the exponential approximation and the effect of varying α.** (**a**) Exact solution vs. exponential approximation. Top: Exact solution and exponential approximation of the detected fluorescence signal as a function of $z$.The inset shows the shift $\Delta z$ between the curves for the range of $z$ from 71 to 78 nm. Bottom: $\Delta z$ as a function of $z$ from the exponential curve ($z_{exp,N_0}$). For $z$ <200 nm, the exponential calculation gives an error in $z$ below 6 nm. In the range of 0-150 nm, the difference in $z$ positions calculated using either of both methods is negligible (<1 nm). (**b**) Influence of α. Top: exponential approximation of the detected fluorescence as a function of $z$ for $\alpha$ in the range of 0.88 to 0.92. This limit correspond to a non-evanescent component of the illumination field 12% and 8% of the total power at $z = 0$, instead of 10% as in our experimental configuration. Variations of $\alpha$ in this range do not introduce distortions greater than 5 nm for $z < 150$ nm.

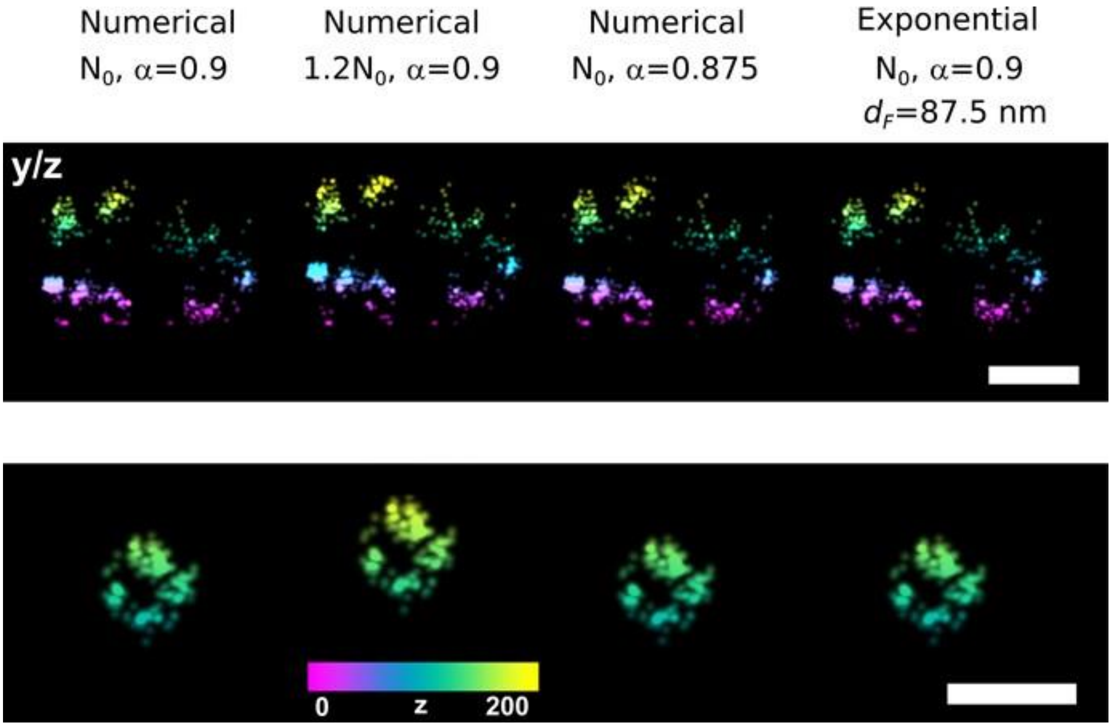

**Supplementary Figure 2. Comparison of side-view reconstructions by SIMPLER using different computation methods and varying α and $N_0$.** Side views (*i.e.* *z*-*y* projections) of a spectrin ring (top) and a microtubule (bottom) obtained with different $z$ -computation approaches. In the first images (left), $z$ was computed numerically using the exact solution using $\alpha = 0.90$ and $N_0 = 50{,}000$. The second images show the influence of choosing another $N_0$ ($1.2N_0$). We can see that the choice of $N_0$ mainly acts as an offset to the $z$-coordinate, influencing principally on the measured distance to the cover-slide. The third column shows the profiles calculated with a different $\alpha$ value. Finally, the side-view reconstructions achieved with the exponential approach ($d_F = 87.5$ nm) are shown in the last column. No significant axial distortions are observed over this range of parameters. Scale bars represent 100 nm.

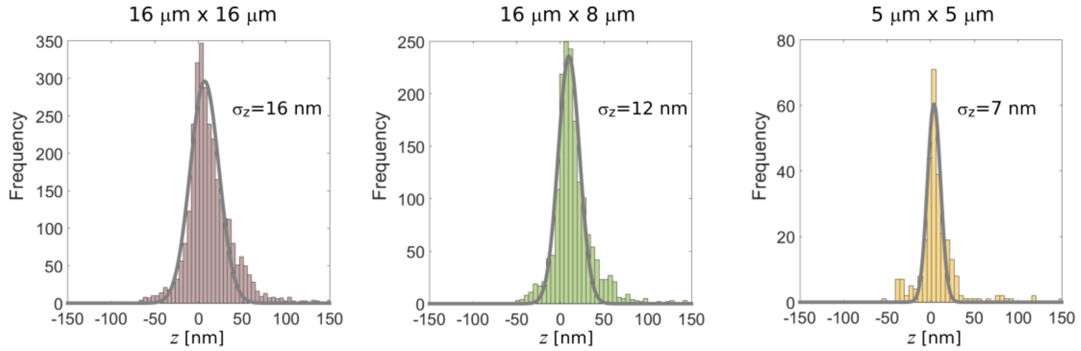

**Supplementary Figure 3. Characterization of the uncertainty in z due to the correction of N by the excitation profile.** Distribution of *z* obtained with SIMPLER – DNA-PAINT for a sample of DNA-Fab fragments adsorbed to the coverslip (z ~ 5 nm). From left to right: the uncertainty in *z* becomes smaller when the area of the region analyzed decreases.

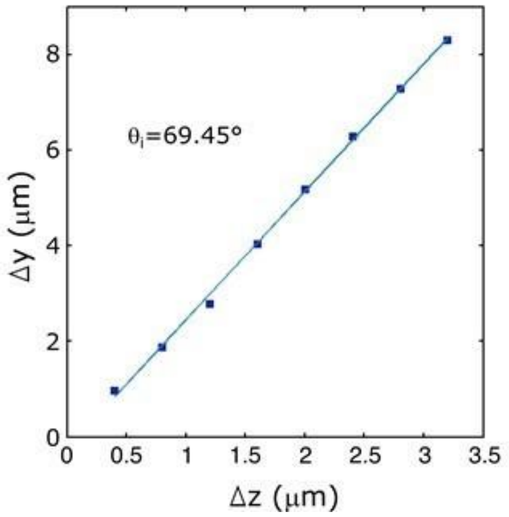

**Supplementary Figure 4. Calibration of the TIRF excitation angle.** Displacement of the excitation laser beam center Δ*y* versus the axial position of the sample Δ*z* (according to Supplementary Method 1). Linear fitting allows determination of $\theta_i$ = 69.45º, in line to the specified limit of the objective used while imaging in the super-resolution microscope set-up 1 (69.5º for an Olympus PlanApo 60x / NA 1.42).

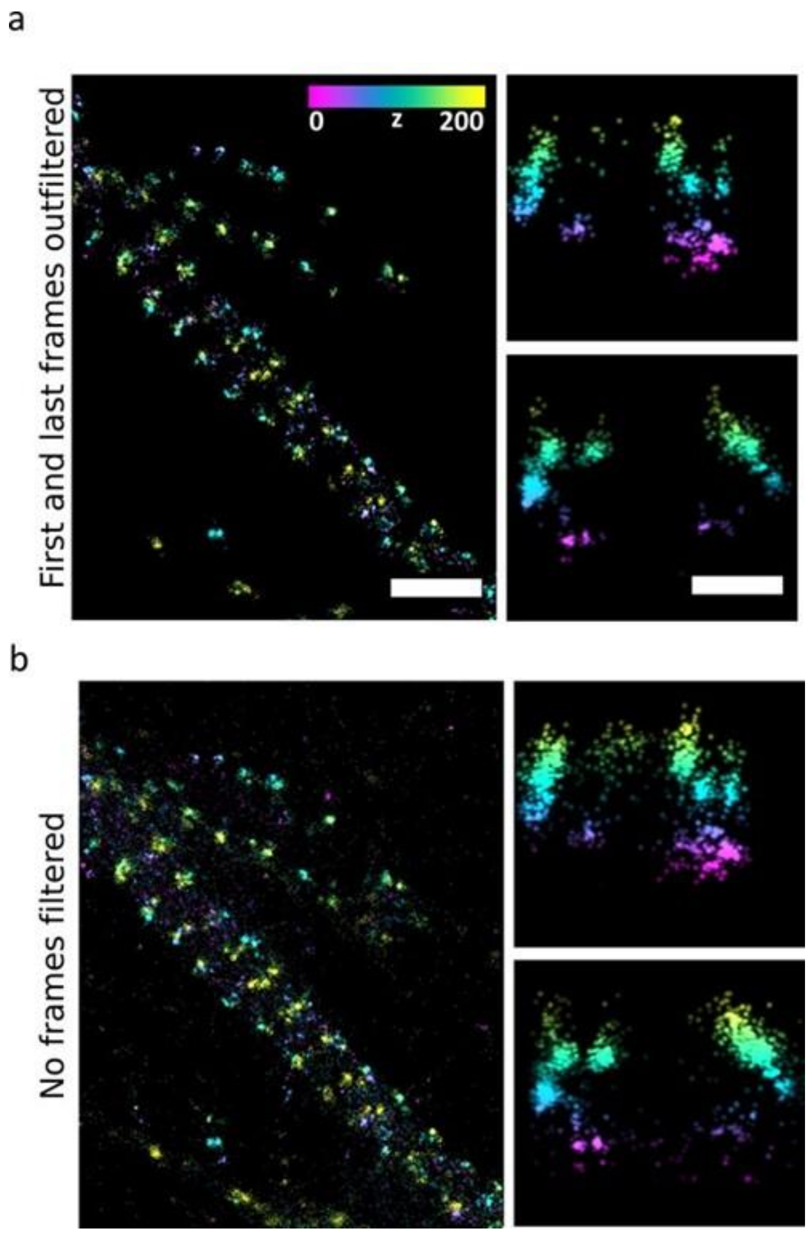

**Supplementary Figure 5. Influence of the first and last frame filtering step on image quality for SIMPLER combined with DNA-PAINT.** Overview image of β2-spectrin rings in neurons and magnified side-view reconstructions, *i.e.* *z*–*y* projections, of the boxed regions in the *x*–*y* view where the rendering was done with (**a**) and without (**b**) the frame filtering step of the localizations (described in Methods and Supplementary Fig. 1). In the *x*–*y* view, the filter's action resembles the one of a density filter, improving contrast by suppressing isolated or unspecific events. In the *z*–*y* projections, we see that the filter suppresses localizations that are wrongly assigned with higher z coordinate due to the incorrectly determined lower photon count. Scale bars represent 1 μm (top view) and 100 nm (side view).

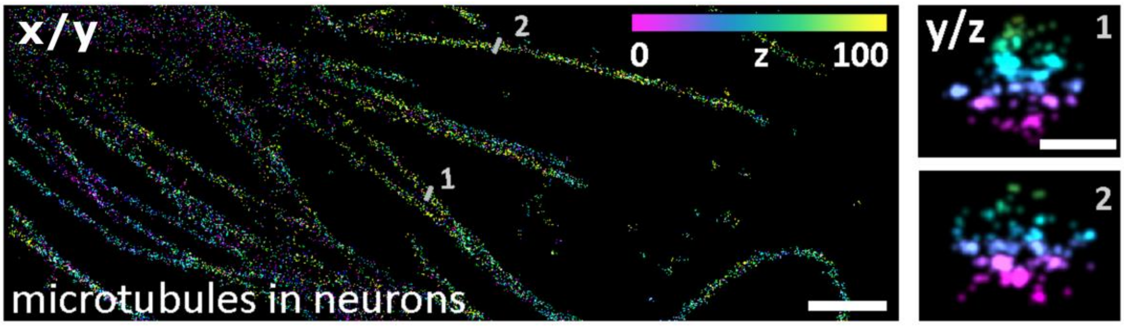

**Supplementary Figure 6. Microtubules from hippocampal neurons immunolabeled for DNA-PAINT super-resolved in 3D using SIMPLER.** Left: top view. Right: magnified side-views along the numbered line in the top view. Scale bars represent 1 μm (top view) and 50 nm (side view).

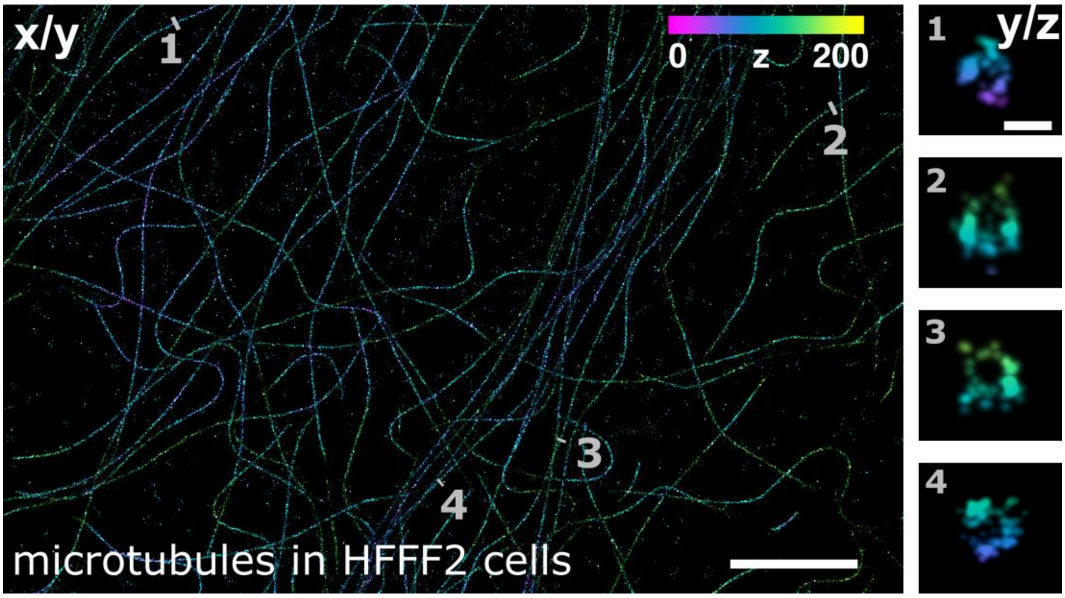

**Supplementary Figure 7. Microtubules from Human Fetal Foreskin Fibroblasts cells, immunolabeled for DNA-PAINT super-resolved in 3D using SIMPLER using a commercial setup (Nikon STORM 5.0).** Left: top view. Right: magnified side-views along the numbered line in the top view. Scale bars represent 4 μm (top view) and 50 nm (side view).

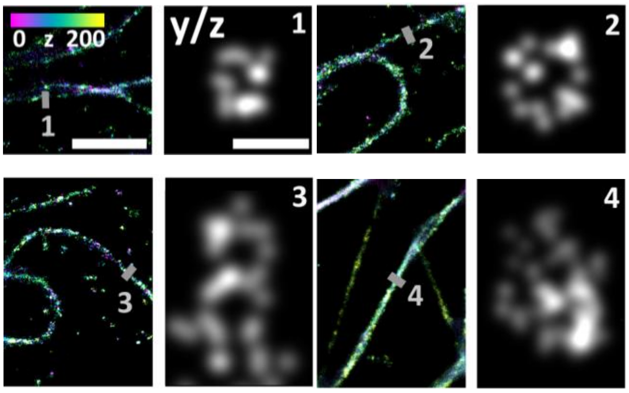

**Supplementary Figure 8. Microtubules immunolabeled for dSTORM super-resolved in 3D using SIMPLER.** (**a**) COS-7 cells (1 to 3) and hippocampal neurons (4). Color-coded: top view. Grayscale: magnified side-views along the numbered line in the top view. Scale bars represent 1 μm (top view) and 50 nm (side view). A $N_0$ value of 6,000 photons was determined.

### Supplementary Tables

| z [nm] | | λ = 500 nm | | | | | λ = 530 | | | |
|---|---|---|---|---|---|---|---|---|---|---|
| | NA | 1.4 | 1.42 | 1.45 | 1.49 | | 1.4 | 1.42 | 1.45 | 1.49 |
| 5 | | 0.593 | 0.628 | 0.680 | 0.727 | | 0.593 | 0.628 | 0.681 | 0.728 |
| 50 | | 0.531 | 0.550 | 0.577 | 0.600 | | 0.534 | 0.554 | 0.583 | 0.606 |
| 100 | | 0.486 | 0.497 | 0.510 | 0.519 | | 0.491 | 0.502 | 0.516 | 0.527 |
| 150 | | 0.454 | 0.460 | 0.466 | 0.470 | | 0.459 | 0.465 | 0.473 | 0.477 |
| 200 | | 0.432 | 0.435 | 0.438 | 0.440 | | 0.437 | 0.440 | 0.444 | 0.446 |
| 250 | | 0.417 | 0.418 | 0.420 | 0.421 | | 0.421 | 0.423 | 0.425 | 0.426 |
| 300 | | 0.409 | 0.410 | 0.410 | 0.411 | | 0.412 | 0.413 | 0.414 | 0.415 |
| 350 | | 0.403 | 0.403 | 0.403 | 0.403 | | 0.403 | 0.403 | 0.404 | 0.404 |
| 400 | | 0.394 | 0.394 | 0.394 | 0.395 | | 0.396 | 0.397 | 0.397 | 0.397 |
| 450 | | 0.391 | 0.391 | 0.391 | 0.391 | | 0.393 | 0.394 | 0.394 | 0.394 |
| 500 | | 0.389 | 0.389 | 0.389 | 0.389 | | 0.392 | 0.392 | 0.392 | 0.392 |
| z [nm] | | λ = 560 | | | | | λ = 590 | | | |
| | NA | 1.4 | 1.42 | 1.45 | 1.49 | | 1.4 | 1.42 | 1.45 | 1.49 |
| 5 | | 0.593 | 0.629 | 0.682 | 0.729 | | 0.594 | 0.630 | 0.683 | 0.730 |
| 50 | | 0.537 | 0.558 | 0.588 | 0.612 | | 0.540 | 0.562 | 0.593 | 0.618 |
| 100 | | 0.495 | 0.507 | 0.523 | 0.535 | | 0.499 | 0.512 | 0.529 | 0.542 |
| 150 | | 0.464 | 0.471 | 0.479 | 0.485 | | 0.469 | 0.476 | 0.485 | 0.492 |
| 200 | | 0.441 | 0.445 | 0.450 | 0.452 | | 0.446 | 0.450 | 0.455 | 0.458 |
| 250 | | 0.425 | 0.427 | 0.429 | 0.431 | | 0.429 | 0.431 | 0.434 | 0.436 |
| 300 | | 0.416 | 0.417 | 0.418 | 0.419 | | 0.419 | 0.421 | 0.422 | 0.423 |
| 350 | | 0.406 | 0.407 | 0.407 | 0.407 | | 0.410 | 0.410 | 0.411 | 0.412 |
| 400 | | 0.400 | 0.400 | 0.400 | 0.401 | | 0.402 | 0.403 | 0.403 | 0.403 |
| 450 | | 0.396 | 0.396 | 0.396 | 0.396 | | 0.398 | 0.398 | 0.398 | 0.399 |
| 500 | | 0.391 | 0.392 | 0.392 | 0.392 | | 0.394 | 0.394 | 0.394 | 0.394 |
| z [nm] | | λ = 620 | | | | | λ = 670 | | | |
| | NA | 1.4 | 1.42 | 1.45 | 1.49 | | 1.4 | 1.42 | 1.45 | 1.49 |
| 5 | | 0.594 | 0.630 | 0.684 | 0.731 | | 0.594 | 0.630 | 0.684 | 0.732 |
| 50 | | 0.543 | 0.565 | 0.597 | 0.624 | | 0.546 | 0.570 | 0.603 | 0.631 |
| 100 | | 0.503 | 0.517 | 0.535 | 0.549 | | 0.509 | 0.524 | 0.543 | 0.559 |
| 150 | | 0.473 | 0.481 | 0.491 | 0.498 | | 0.479 | 0.488 | 0.500 | 0.508 |
| 200 | | 0.450 | 0.455 | 0.460 | 0.464 | | 0.456 | 0.462 | 0.469 | 0.473 |
| 250 | | 0.433 | 0.436 | 0.439 | 0.441 | | 0.439 | 0.442 | 0.446 | 0.449 |
| 300 | | 0.423 | 0.425 | 0.426 | 0.427 | | 0.428 | 0.431 | 0.433 | 0.434 |
| 350 | | 0.413 | 0.414 | 0.415 | 0.415 | | 0.418 | 0.419 | 0.420 | 0.421 |
| 400 | | 0.405 | 0.406 | 0.407 | 0.407 | | 0.409 | 0.410 | 0.411 | 0.412 |
| 450 | | 0.401 | 0.401 | 0.401 | 0.402 | | 0.406 | 0.406 | 0.407 | 0.407 |
| 500 | | 0.396 | 0.396 | 0.396 | 0.397 | | 0.400 | 0.400 | 0.400 | 0.401 |
| | | λ = 700 | | | | | λ = 720 | | | |
| z [nm] | NA | 1.4 | 1.42 | 1.45 | 1.49 | | 1.4 | 1.42 | 1.45 | 1.49 |
| 5 | | 0.598 | 0.634 | 0.688 | 0.735 | | 0.593 | 0.630 | 0.684 | 0.732 |
| 50 | | 0.546 | 0.570 | 0.604 | 0.632 | | 0.549 | 0.574 | 0.609 | 0.638 |
| 100 | | 0.509 | 0.524 | 0.544 | 0.560 | | 0.514 | 0.530 | 0.552 | 0.569 |
| 150 | | 0.479 | 0.488 | 0.500 | 0.509 | | 0.485 | 0.495 | 0.509 | 0.518 |
| 200 | | 0.456 | 0.462 | 0.469 | 0.474 | | 0.463 | 0.469 | 0.478 | 0.483 |
| 250 | | 0.438 | 0.442 | 0.446 | 0.449 | | 0.445 | 0.449 | 0.454 | 0.458 |
| 300 | | 0.428 | 0.430 | 0.433 | 0.434 | | 0.434 | 0.437 | 0.440 | 0.442 |
| 350 | | 0.417 | 0.418 | 0.420 | 0.421 | | 0.423 | 0.425 | 0.427 | 0.428 |
| 400 | | 0.409 | 0.410 | 0.411 | 0.411 | | 0.414 | 0.416 | 0.417 | 0.417 |
| 450 | | 0.405 | 0.405 | 0.406 | 0.406 | | 0.410 | 0.411 | 0.411 | 0.412 |
| 500 | | 0.402 | 0.402 | 0.403 | 0.403 | | 0.404 | 0.404 | 0.405 | 0.405 |

**Supplementary Table 1. Axial dependence of the collected fluorescence signal.** Values of $CF_{avg}$ as a function of $z$ for various values of maximum emission wavelength ($\lambda$) and numerical aperture (NA), which together with the axial dependence of the illumination field allow users to extract the decay ($d_F$) and the background constant ($\alpha_F$). Alternatively, users can directly obtain $d_F$ and $\alpha_F$ by simply input of their experimental parameters in the supplemented MATLAB script (Supplementary Software 1).

### Supplementary Methods

#### Supplementary Method 1: TIRF angle calibration

Incident angle of the excitation beam was determined as previously described.[3] Briefly, 1 µM AlexaFluor 647 solution was illuminated with an incident angle $\theta_i > \theta_c$ and the sample was translated in *z*-direction from $z = 0$ to $z = 10$ µm, in 0.4 µm steps (Prior ProScan III). As a consequence of the *z*-translation of the sample the excitation spot was displaced in a lateral direction (*y*). The value of $\theta_i$ was obtained by fitting the dependence of lateral movement of the center of the excitation beam on the *z*-translation with a linear regression, where $\Delta y = m\Delta z + c$ and $\arctan(m) = \theta_i$ (See calibration data and fit in Supplementary Fig. 4).

#### Supplementary Method 2: DNA-antibody coupling reaction

DNA labelling of a fragment secondary antibody (donkey anti-mouse IgM, 715-007-003 or donkey anti-rabbit IgM, 711-007-003, Jackson ImmunoResearch) was performed using the maleimidePEG2-succinimidyl ester coupling reaction.[4] In order to reduce the thiolated DNA for the maleimide reaction, 15 µL of 1 mM thiol-DNA (5'-TATGTAACTTT-3'-Thiol and 5'-ATTACTTCTTT-3'-Thiol, biomers.net GmbH, for the donkey anti-mouse and the donkey anti-rabbit conjugates respectively) were incubated with 35 µL of 250 mM DDT (Thermo Fisher Scientific) freshly prepared solution (1.5 mM EDTA, 0.5x PBS, pH 7.2) on a shaker, in the dark, for 2 h at room temperature. 30 min after the reduction of the thiol-DNA started, 30 µL of 26 µM fragment antibody was incubated with 0.7 µL of 23.5 mM Maleimide-PEG2-succinimidyl ester (Sigma-Aldrich) solution on a shaker, in the dark, for 90 min at 4ºC. Prior DNA-antibody conjugation, both sets of reactions were purified using an illustra MicroSpin G-25 column (GE Healthcare) to remove excess of DDT and a Zeba desalting column (Thermo Fisher Scientific) to remove excess of cross-linker. Next, both flow-through of the columns were mixed and incubated on a shaker, in the dark, overnight at 4ºC. The next day, DNA excess was removed by Amicon spin filtration (30 kDa). Antibody-DNA concentration was measure with the NanoDrop spectrophotometer and adjusted to 14 µM with PBS. DNA-labelled antibodies were store for a maximum of 6 months at 4 °C.
