## Supplementary Software for "Three-dimensional total-internal reflection fluorescence nanoscopy with nanometric axial resolution by photometric localization of single molecules": getParameters_SIMPLE.m

```
function [dF, alphaF] = getParameters_SIMPLE(lambda_exc,NA,lambda_em,angle,nS,nI,alpha);

dF=[];
alphaF=[];
dif=[];

if angle==0
    angle = asin(NA/nI);
else 
    angle=deg2rad(angle);
end

if alpha==0
    alpha = 0.9;
else 
    alpha=alpha;
end

z =[5 50 100 150 200 250 300 350 400 450 500];
z_fit=5:0.5:500;
lambda_em_max =[500 530 560 590 620 670 700 720];
[dif,i_lambda_em]= min(abs(lambda_em_max-lambda_em));

%Axial dependece of the excitation field

d = lambda_exc/(4*pi()*sqrt(nI^2*(sin(angle)^2)-nS^2));
I_exc = alpha*exp(-z_fit./d)+(1-alpha); 

    
% Axial dependece of the fraction of fluorescence collected by a microscope 
% objective

if NA==1.42
    load DF_NA1.42.txt;    
    DFi = DF_NA1_42(:,i_lambda_em);

elseif NA==1.45;        
     load DF_NA1.45.txt;
     DFi = DF_NA1_45(:,i_lambda_em);
     
     elseif NA==1.40;        
     load DF_NA1.40.txt;
     DFi = DF_NA1_40(:,i_lambda_em);
     
elseif NA==1.49;
         load DF_NA1.49.txt;
         DFi = DF_NA1_49(:,i_lambda_em);
end

DFi_interp=interp1(z,DFi,z_fit);
I_total = I_exc.*DFi_interp;


% Fit F to "F = alphaF*exp(-z/dF)+(1-alphaF)"

f = @(b,z_fit) b(1).*exp(-b(2).*z_fit)+b(3);                                     % Objective Function
B = fminsearch(@(b) norm(I_total - f(b,z_fit)), [0.6; 0.01; 0.05]);                  % Estimate Parameters
z_fit_from0 = 0:0.5:500;
figure
plot(z_fit, I_total, 'pg')
hold on
plot(z_fit, f(B,z_fit), '-r')
hold off
grid
xlabel('z')
ylabel('F')
title('F and fit vs. z');
text(27, 105, sprintf('f(x) = %.1f\\cdote^{%.3f\\cdotx}%+.1f', B))
figure
plot (z_fit_from0,(f(B,z_fit_from0)/(B(1,1)+B(3,1))),'-r')
grid
xlabel('z')
ylabel('F')
title('Normalized fit vs. z');
text(27, 105, sprintf('f(x) = %.1f\\cdote^{%.3f\\cdotx}%+.1f', B))
alphaF = (B(1,1)/(B(1,1)+B(3,1)))
dF = 1/B(2,1)
end
     
     
```
