## Supplementary Software for "Three-dimensional total-internal reflection fluorescence nanoscopy with nanometric axial resolution by photometric localization of single molecules": SIMPLE.m

```
%% SIMPLE

%   This code uses the experimental conditions of your single-molecule 
%   based super resolution experiments, including microscope set-up 
%   (NA, incident TIR angle, evanescent component,laser)and sample 
%   conditions (fluorophore) to provide you with two parameters:
%  
%       - decay (dF)
%       - background constant (alphaF) 
%
%   Once dF and alphaF has been calculated, you can use them to decode z 
%   information with the exponential approximation,
%   
%   z = dF*log{(alphaF*N0)/[N-(1-alphaF)*N0)]}
%
%   where N corresponds to the the number of detected photons per frame,
%   and N0 is the number of detected photons per frame at z = 0.

%% Answer the following questions to obtain dF and alphaF:

clear all; 

% 1. Which is the excitation wavelength? 
%    Typical lasers lines include: 480 nm, 532 nm, 561 nm and 642 nm.  
    
        lambda_exc = 642;
        
% 2. Which is the maximun emission wavelength of your fluorophore? 
%    Alternatively, you can also input the center of your emission filter

        lambda_em = 700;

% 3. Which is the NA of the TIR objective?
%    Choose between typical NA values of 1.42, 1.45, 1.49.
       
        NA = 1.40;

% 4. Which is the refractive index (nI) of the medium of incidence? (check the
%    oil immersion objetive nI)
%
        nI = 1.517; 
%
% 5. Which is your angle of incidence (in degrees)?
%    If this value is not input (i.e. angle = 0), we will use the  maximum angle for your 
%    objective. 

        angle = 69.45;

% 6. Which is the relative contribution of the non-evanescent field?
%    This value usually represent a 10% of the total power at z = 0 for most 
%    objective-type TIR configurations. If this value is not input (i.e. 
%    alpha = 0) we will use (1-alpha) = 0.1.
     
        alpha = 0.9;
     
% 7. Which is the refraction medium of your sample? 
%    Typical biological samples are in water and hence have a refractive 
%    index of 1.33

        nS = 1.33; 

[dF, alphaF] = getParameters_SIMPLE(lambda_exc,NA,lambda_em,angle,nS,nI,alpha);
```
